# Metagenomics reveals diet-specific specialization of bacterial communities in fungus gardens of grass- and dicot-cutter ants

**DOI:** 10.1101/250993

**Authors:** Lily Khadempour, Huan Fan, Ken Keefover-Ring, Camila Carlos, Nilson S. Nagamoto, Miranda A. Dam, Monica T. Pupo, Cameron R. Currie

**Affiliations:** Department of Bacteriology, University of Wisconsin-Madison, Madison, WI, USA, 53706; Department of Integrative Biology, University of Wisconsin-Madison, Madison, WI, USA, 53706; Department of Energy Great Lakes Bioenergy Research Center, University of Wisconsin-Madison, Madison, WI, USA, 53706; Departments of Botany and Geography, University of Wisconsin-Madison, Madison, WI, USA, 53706; Department of Plant Protection, UNESP – São Paulo State University, Botucatu, SP, Brazil, CEP: 18603-970; Department of Nutritional Sciences, University of Wisconsin-Madison, Madison, WI, USA 53706; School of Pharmaceutical Sciences of Ribeirão Preto, University of São Paulo, Ribeirão Preto, São Paulo, Brazil, CEP: 14040-903

## Abstract

Leaf-cutter ants in the genus *Atta* are dominant herbivores in the Neotropics. While most species of *Atta* cut dicots to incorporate into their fungus gardens, some species specialize on grasses. Here we examine the bacterial community associated with the fungus gardens of grass- and dicot-cutter ants to examine how changes in substrate input affect the bacterial community. We sequenced the metagenomes of 12 *Atta* fungus gardens, across four species of ants, with a total of 5.316 Gbp of sequence data. We show significant differences in the fungus garden bacterial community composition between dicot- and grass-cutter ants, with grass-cutter ants having lower diversity. Reflecting this difference in community composition, the bacterial functional profiles between the fungus gardens are significantly different. Specifically, grass-cutter ant fungus garden metagenomes are particularly enriched for genes responsible for amino acid, siderophore, and terpenoid biosynthesis while dicot-cutter ant fungus gardens metagenomes are enriched in genes involved in membrane transport. These differences in bacterial community composition and functional capacity show that different substrate inputs matter for fungus garden bacteria, and sheds light on the potential role of bacteria in mediating the ants’ transition to the use of a novel substrate.

## Introduction

Understanding the role of microbial symbionts in aiding nutrient acquisition is fundamental to understanding the biology of herbivores. Herbivores are the most diverse and abundant animals on earth, and play an important role ecosystem functioning and nutrient cycling (Ricklefs and Miller, 2000; Huntly, 1991). Most herbivores host microbial symbionts that serve as an interface between them and the plants that they consume. These microbes can compensate for the hosts’ lack of physiological capacity to obtain energy and nutrients from plants (Hansen and Moran, 2013). Herbivore microbial symbionts, often residing in the guts of animals, have been implicated in aiding plant biomass breakdown (Hess *et al.*, 2011; Kudo, 2009; Talbot, 1977; Adams *et al.*, 2011), plant defense compound remediation (Wang *et al.*, 2010; Adams *et al.*, 2013; Boone *et al.*, 2013), and nutrient supplementation (Warnecke *et al.*, 2007; Hansen and Moran, 2011; LeBlanc *et al.*, 2013). Microbial communities differ between hosts that specialize on different substrates (Muegge *et al.*, 2011) and changes in these communities and their functional capacity are integral to their hosts’ transition to utilizing novel substrates (Hammer and Bowers, 2015; Delsuc *et al.*, 2013; Li *et al.*, 2015; Kohl *et al.*, 2014; 2016). This phenomenon is not isolated to herbivores – in many systems, gut microbial communities are influence by the inputs of their hosts (Goffredi *et al.*, 2005; Roman *et al.*, 2015; Muegge *et al.*, 2011; Youngblut *et al.*, 2019; Wang *et al.*, 2011; Li *et al.*, 2016).

Leaf-cutter ants represent a paradigmatic example of the microbial mediation of herbivory. They are dominant herbivores in the Neotropics, consuming up to an estimated 17% of foliar biomass in the systems in which they live (Costa *et al.*, 2008). These ants have a significant impact on their surrounding ecosystems, due to the volume of plant biomass they consume and soil that they excavate in building their underground colonies (Herz *et al.*, 2007; Costa *et al.*, 2008; Fowler *et al.*, 1986; Moutinho *et al.*, 2003; Gutiérrez and Jones, 2006). Leaf-cutter ants lack the capacity to break down recalcitrant plant material. Instead, they gain access to the nutrients in plant biomass by farming a fungus, *Leucoagaricus gongylophorus*, which enzymatically breaks down recalcitrant biomass in the leaf material that the ants forage (Aylward *et al.*, 2013; Khadempour *et al.*, 2016; Suen *et al.*, 2011a; Grell *et al.*, 2013; Kooij *et al.*, 2011; Nagamoto *et al.*, 2011). *Leucoagaricus gongylophorus* produces gongylidia, specialized hyphal swellings that contain an abundance of sugars and lipids, that the ants consume and feed to larvae (Bass and Cherrett, 1995; North *et al.*, 1997). In the leaf-cutter ant system, the fungus garden serves as the ants’ external gut, and the combination of ants and fungus garden can be considered a holobiont. While the ants primarily consume fungus, we consider a leaf-cutter ant colony as an individual holobiont that interacts with its environment as a herbivore (Hussa and Goodrich-Blair, 2013; Bordenstein and Theis, 2015).

Recent work has revealed that a community of bacteria reside within leaf-cutter ant fungus gardens (Suen *et al.*, 2010; Aylward *et al.*, 2012; Moreira-Soto *et al.*, 2017). These communities are dominated by Gammaproteobacteria, and consistently contained strains of *Pseudomonas, Enterobacter* and either *Rahnella* or *Pantoea,* and are highly similar to communities of bacteria associated with other fungus-farming insects (Aylward *et al.*, 2014; 2012; Suen *et al.*, 2011a). Some garden bacteria are vertically transmitted, maternally through the fungus pellets that alate queens use to establish new fungus gardens (Moreira-Soto *et al.*, 2017). The community consistency and their vertical transmission, suggest that the bacterial communities are important to the fitness of their hosts. One study, by Pinto-Tomás *et al*. (2009) showed that *Pantoea* and *Klebsiella* bacteria that are found in leaf-cutter ant fungus gardens fix nitrogen that supplements the ant diet, which is important for a strict herbivorous system. Nevertheless, the functional role of most garden bacteria remains unknown.

While most leaf-cutter ants use dicots, three species of *Atta* are specialized on cutting grass, and another three species cut both grasses and dicots (Fowler *et al.*, 1986). All previous studies on the microbial community in leaf-cutter ant fungus gardens have been focused on dicot-cutting ants, likely because dicot-cutters are more common and grass-cutter ants are notoriously difficult to maintain in the lab (Nagamoto *et al.*, 2009). In this study, we compare the bacterial communities of fungus gardens from ants that cut grass and dicots. Given that grasses and dicots differ in terms of the cell wall composition (Popper and Tuohy, 2010; Ding and Himmel, 2008), plant defense compounds (Wetterer, 1994; Mariaca *et al.*, 1997) and nutrient availability (Mattson, 1980; Winkler and Herbst, 2004), we hypothesize that the bacterial community in these fungus gardens will differ in terms of community composition and functional capacity, in response to the different composition of the substrates the ants incorporate into their gardens. To address this, we collected fungus gardens from grass- and dicot-cutter ants and obtained their bacterial community metagenomes using Illumina sequencing. We analyzed the bacterial community in terms of its taxonomic composition and its functional capacity. We also conducted analyses on the fungus gardens to determine their plant composition, their nutritional composition and their plant defense compound contents.

## Methods

### Collection of fungus garden

Fungus gardens were collected on the campuses of the University of São Paulo (USP) in Ribeirão Preto, SP, Brazil and the São Paulo State University (UNESP) in Botucatu, SP, Brazil. Collection dates and GPS coordinates are listed in Table 1. We collected fungus gardens from four species of *Atta* leaf-cutter ants: *A. bisphaerica* and *A. capiguara*, which are both described as grass-cutters, *A. laevigata*, which is described as a grass and dicot-cutter, and *A. sexdens*, which is described as a dicot-cutter (Fowler *et al.*, 1986).

**Table 1.**
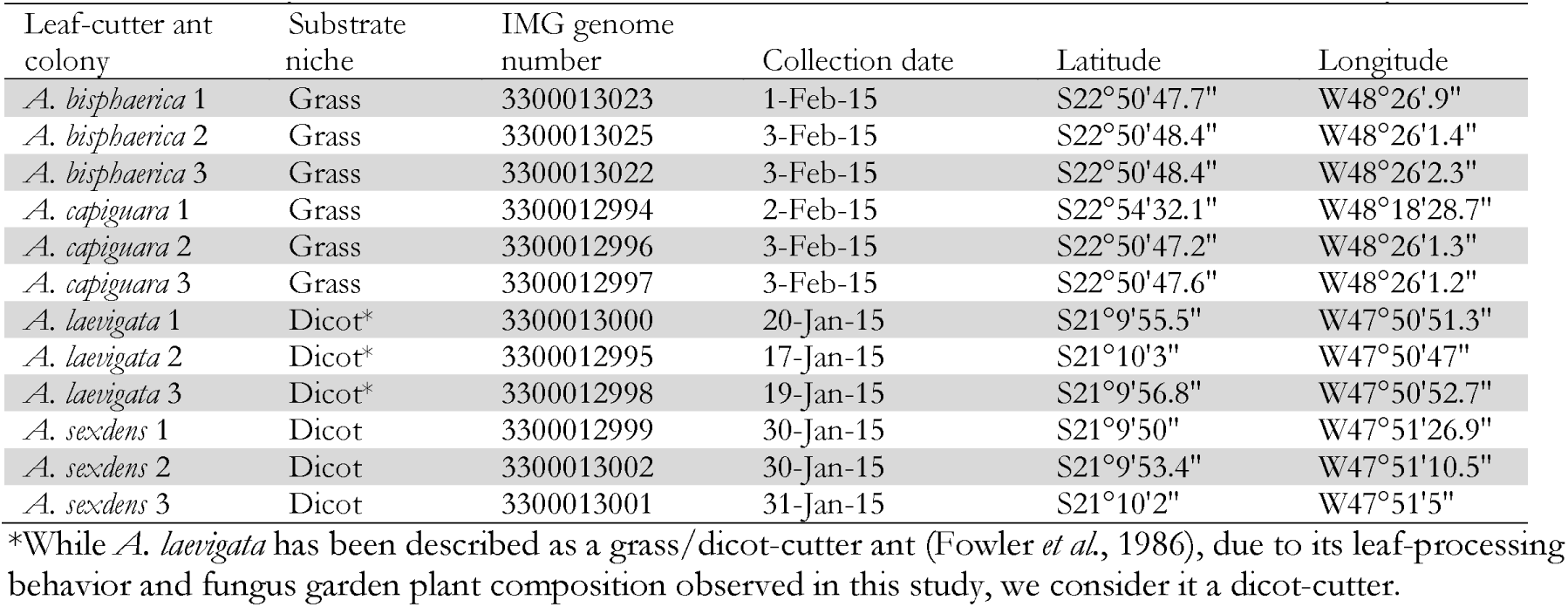
Summary of collection details for leaf-cutter ant colonies used in this study

From each species we collected from three independent colonies. To collect the fungus gardens, we identified the ant species by worker morphology then followed the entrance tunnel by digging until we found a fungus garden. Care was taken to expose fungus garden chambers from the side, to avoid damaging the garden with digging tools and to avoid contamination with surrounding soil. Fungus gardens were transported to the laboratory and aseptically transferred into 50 mL conical tubes. When we sampled multiple chambers from one colony, they were always adjacent to each other and were superficially similar in appearance, texture and odor. The majority of worker ants were removed from the fungus garden material before being transferred to the tubes. In order to further reduce the chance of soil contamination, only intact fungus garden from the central region of the fungal mass was included in the tubes. Once filled, the tubes were frozen in liquid nitrogen and stored at −80°C. At least four 50 mL conical tubes were filled from each colony. For each colony, two tubes were combined and used for metagenomics, one tube was used for gas chromatography, and one tube was used for iron content measurements.

### DNA extraction

To target the bacteria in the fungus gardens, DNA was extracted by first using a differential centrifugation method (Aylward *et al.*, 2012). PBS buffer with 1% Tween 80 was added to the tubes and they were shaken for 30 min on a vortex. They were then kept at 4°C for 30 min so that large particles would settle. The liquid portion was decanted and passed through a 40 *μ*m filter. The filtrate was centrifuged for 30 min at 4°C at 4300 rpm (Beckman Coulter X-14R centrifuge with an SX4750 swinging bucket), after which a bacterial cell pellet was formed and the liquid was removed. This process was repeated with the original fungus garden tube to wash off any remaining bacterial cells from the leaf material, but we did not notice an appreciable difference in the size of the pellet after the second wash. DNA was extracted from the cell pellet using the Qiagen Plant DNA Extraction Maxi Kit (Qiagen, Hilden, Germany). The remaining leaf material from the fungus gardens was photographed after the differential centrifugation, to demonstrate the difference in leaf material consistency (Figure 1).

**Figure 1.**
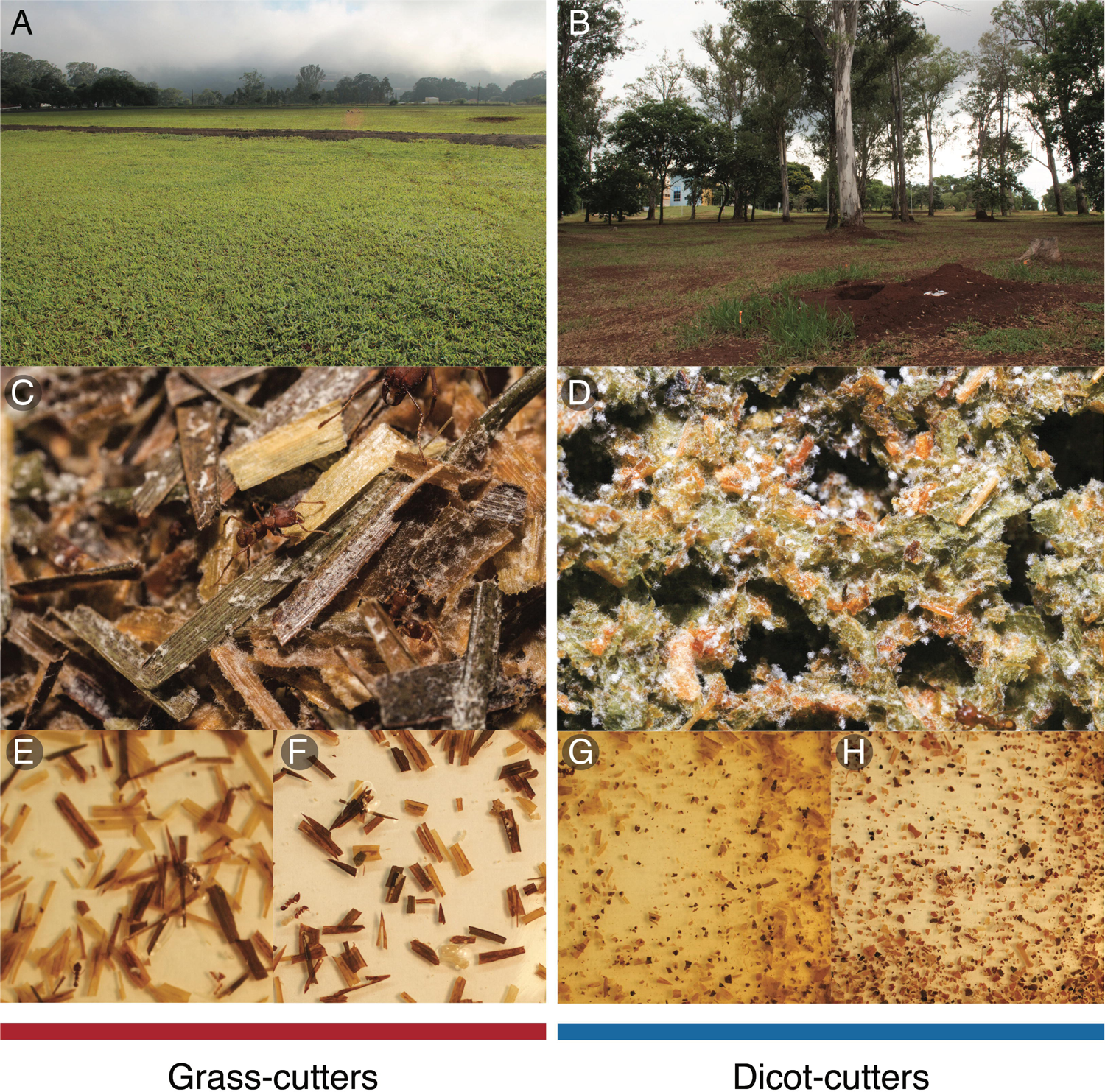
Grass- and dicot-cutter ants differ in the niches that they occupy, and the way that they cut and process leaf material. Field sites in (A) Botucatu, SP and (B) Ribeirão Preto, SP, Brazil. Fungus gardens of (C) grass- and (D) dicot-cutter ants. C. Visual inspection of leaf material from leaf-cutter ant fungus gardens demonstrates the degree of trituration that the different ants complete, with grass-cutters leaving the leaf material more intact (E – *A. bisphaerica* and F – *A. capiguara*), while dicot-cutters triturate to the point of producing unrecognizable leaf fragments (G – *A. laevigata* and H – *A. sexdens*).

### DNA sequencing and assembly

All metagenomic sequencing was conducted at the Joint Genome Institute (JGI) in Walnut Creek, CA. Since some of the DNA concentrations were too low for standard library prep, a low-input prep was completed for all of the samples. For each sample, 10 ng of DNA was sheared to 300 bp using the Covaris LE220 (Covaris) and size selected using SPRI beads (Beckman Coulter). The fragments were treated with end-repair, A-tailing, and ligation of Illumina compatible adapters (IDT, Inc) using the KAPA-Illumina library creation kit (KAPA Biosystems) and 10 cycles of PCR was used to enrich for the final library. The prepared libraries were quantified using KAPA Biosystems next-generation sequencing library qPCR kit and run on a Roche LightCycler 480 real-time PCR instrument. The quantified libraries were then prepared for sequencing on the Illumina HiSeq sequencing platform utilizing a TruSeq Rapid paired-end cluster kit, v4. Sequencing of the flowcell was performed on the Illumina HiSeq2500 sequencer using HiSeq TruSeq SBS sequencing kits, following a 2×150 indexed run recipe. BBDuk adapter trimming (Bushnell, 2017) was used to remove known Illumina adapters. The reads were then processed using BBDuk filtering and trimming. Read ends were trimmed where quality values were less than 12. We discarded read pairs that fit certain criteria: those containing more than three ambiguous bases, or quality scores (before trimming) averaging less than three over the read, or length under 51 bp after trimming, as well as reads matching Illumina artifact, spike-ins or phiX. Trimmed, screened, paired-end Illumina reads were assembled using the megahit assembler using with the “--k-list 23,43,63,83,103,123” option. Functional annotation and taxonomic classification were performed using the Integrated Microbial Genomes (IMG) pipeline (Chen *et al.*, 2018).

### Plant genus richness

To determine the richness of plant substrate integrated in the fungus gardens of the ants, we used JGI’s IMG database “find gene” function to retrieve all genes annotated as *MatK* from the dataset. *MatK* is a widely used chloroplast plant DNA barcode (Hollingsworth *et al.*, 2011). Retrieved *MatK* sequences for each metagenome were identified using BLASTn. The best match for each sequence was identified first to the species level (where all matches were at least 98%) but to ensure consistent and reliable certainty with the identified plants, we identified all sequences to the genus level. Because most of the plant biomass was removed from samples before DNA extraction only presence/absence of genera were considered, not abundance.

### Bacterial taxonomic analysis

Relative abundance of bacterial taxa (classes and genera) were determined using MATAM (Pericard *et al.*, 2018). Reads originated from 16s genes were identified and assembled into contigs, and we set the coverage threshold to be 500x. We also used MATAM for taxonomic assignment using the “perform taxonomic assignment” option. This calls an RDP Classifier (Wang *et al.*, 2007), which is a naive Bayesian classifier with the default training model “16srrna”. MATAM also reports the coverage of each contig to be used for the calculation of relative abundance; here we only kept those with coverage above 1 as a quality control, excluded contigs that could not be identified, and collapsed taxa that represented less than 1% of the community into one category. We used the relative abundances of each phylum and genus to run a non-metric multidimensional analysis (NMDS) using a Bray-Curtis dissimilarity index with the vegan package in the R statistical programming environment (Oksanen *et al.*, 2013; R Core Team, 2013). Relative abundance was calculated as the proportion of each bacterial genus or class compared to the total quantity of bacteria in the sample. Also using the vegan package, we used ANOSIM and PERMANOVA to determine if groups (grass-cutters vs. dicot-cutters) were significantly different, and we used the Shannon diversity index to compare the diversity of each sample on the basis of the bacterial genera present. To test whether specific genera have significantly different relative abundances between grass- and dicot-cutter ant fungus gardens, we used DESeq2 in the R statistical programming environment (Love *et al.*, 2014).

### Bacterial functional analysis

In order to make functional comparisons of the bacteria in grass- and dicot-cutter fungus gardens, we used the Kyoto Encyclopedia of Genes and Genomes (KEGG) annotations of the metagenomes through IMG’s KEGG Orthology (KO) pipeline, which is part of JGI’s standard operating procedure (Huntemann *et al.*, 2016). Briefly, genes were associated with KO terms (Kanehisa *et al.*, 2014) based on USEARCH 6.0.294 results (Edgar, 2010) and were filtered for minimum identity matches and gene sequence coverage. For an overall comparison of functional differences between the fungus gardens, we used the same ordination and statistical methods as for bacterial genus relative abundance. As with genus group differences, we used DESeq2 to determine what genes are significantly enriched between grass- and dicot-cutter ant fungus gardens. Since DESeq2 requires inputs to be integers, we used number of gene copies per million genes in the metagenomes as our input (Alneberg *et al.*, 2014).

### Iron content

Separate 50 mL tubes of fungus garden material, from the same colonies as above, were used for determination of iron content. All ants were removed from fungus gardens then the remaining fungus garden material (triturated plants covered in fungus) was analyzed at the UW Soil and Forage Lab in Marshfield, WI, using standard methods. Approximately 0.5 g of dried and ground fungus garden material was weighed out into a folin digestion tube. The material was then digested in 5 mL of concentrated nitric acid (67-70%), and heated to 120°C for four hours. Next, 1 mL of hydrogen peroxide (30%) was added, and the samples were heated for a further 20 min before being diluted and analyzed by inductively coupled plasma optical emission spectroscopy (ICP-OES) (Fassel and Kniseley, 1974).

## Results

### Metagenomic statistics

A summary of metagenome statistics is presented in (Table 2). A total of 5.316 Gbp of assembled sequence data was produced in this study, with an average of 443 Mbp per metagenome. The smallest metagenome was from the grass-cutter colony *A. capiguara* 1 at 148.7 Mbp, and the largest metagenome was from the dicot-cutter colony *A. sexdens* 2 at 812.9 Mbp. Maximum scaffold lengths ranged from 61.96 Kbp to 701.42 Kbp, with an average maximum scaffold length of 266.6 Kbp. Between 91.63% and 99.31% of reads were aligned.

**Table 2.**
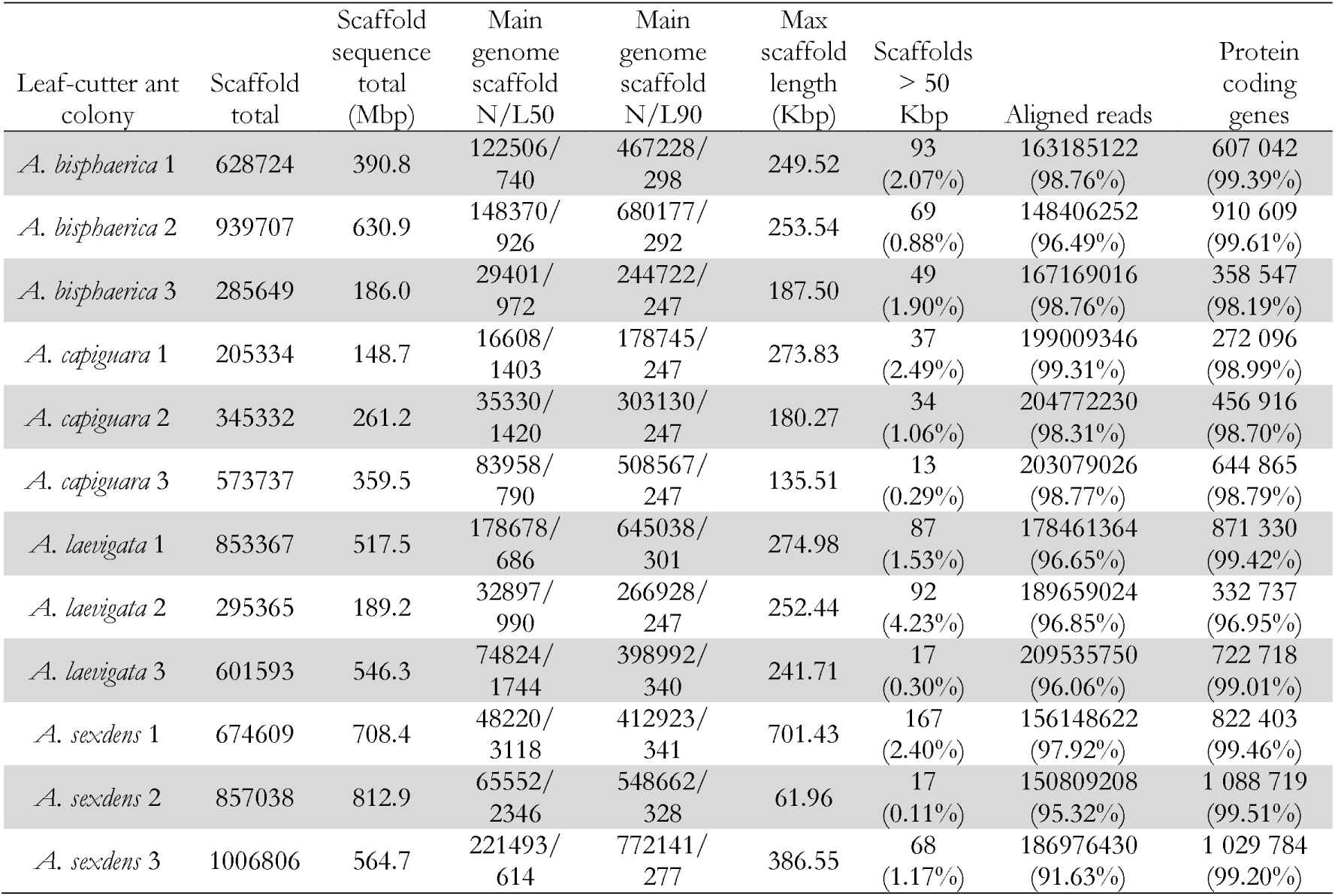
Metagenome sequencing statistics for leaf-cutter ant fungus gardens

### Bacterial taxonomic analysis

Overall, *Atta* spp. fungus garden bacterial communities differed from each other, depending on if the ants were grass- or dicot-cutters (Figure 2). All fungus garden bacterial communities were dominated by Proteobacteria. Grass-cutter ant fungus gardens were comprised mostly of Gammaproteobacteria (94%), and while this their proportion was still high, it was lower in dicot-cutter ant fungus gardens (54%). Alphaproteobacteria were also a dominant class in dicot-cutter ant fungus gardens where they were 34% of the average bacterial community. The most relatively abundant genus in all the fungus gardens was *Pantoea*, at 67% and 16% of the grass- and dicot-cutter ant fungus gardens, respectively. Other abundant genera were *Pseudomonas* at 18% and 16% of the grass- and dicot-cutter ant fungus gardens, respectively, and *Gluconobacter*, which was 19% of the bacterial community in dicot-cutter ant fungus gardens, but was not found at all in grass-cutter ant fungus gardens. *Burkholderia* and *Enterobacter* were found in both fungus gardens at an average proportion of 2% of the bacterial community (Figure 3A). Both grass- and dicot-cutter ant fungus gardens contained bacterial genera that were unique to the their respective groups, but there were more of these substrate-specific bacteria in the dicot-cutter ant fungus gardens, and this contributed to the higher overall diversity of bacteria in the dicot-cutter ant fungus gardens (Shannon diversity index of 1.46) compared to the grass-cutter ant fungus gardens (Shannon diversity index of 0.71) (Figure 3B). This is also reflected in the mean genus richness, which were 7.7 and 4.3 genera in the dicot- and grass-cutter ant fungus gardens, respectively (Figure 3C). Despite these patterns, DESeq2 analysis did not find significant differences between particular high-abundant genera between the two types of fungus gardens, likely due to low power in the analysis.

**Figure 2.**
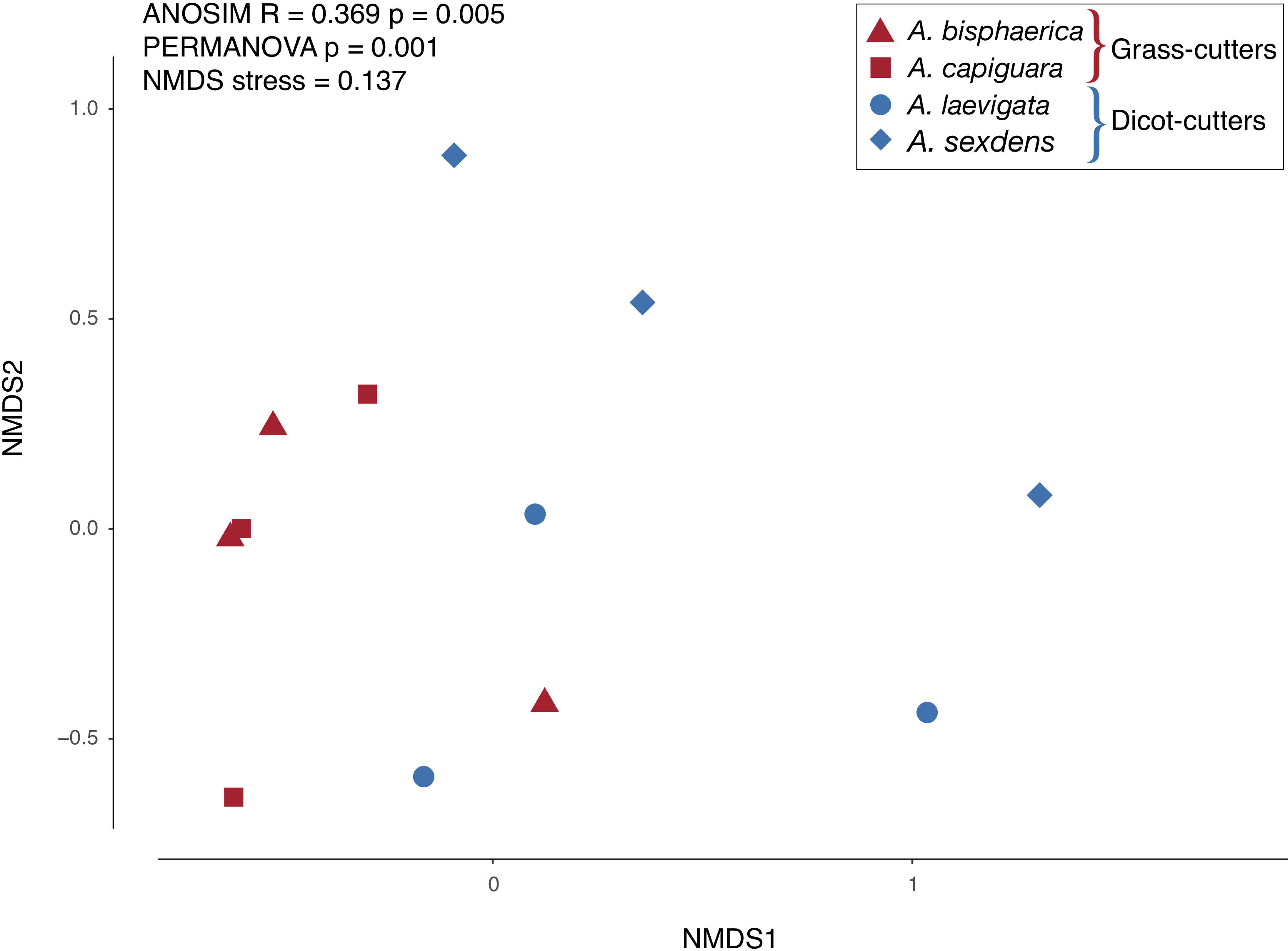
NMDS plot of the relative abundance of bacterial genera in fungus gardens of grass- and dicot-cutter ants. Grass- and dicot-cutter fungus garden bacterial communities are significantly different.

**Figure 3.**
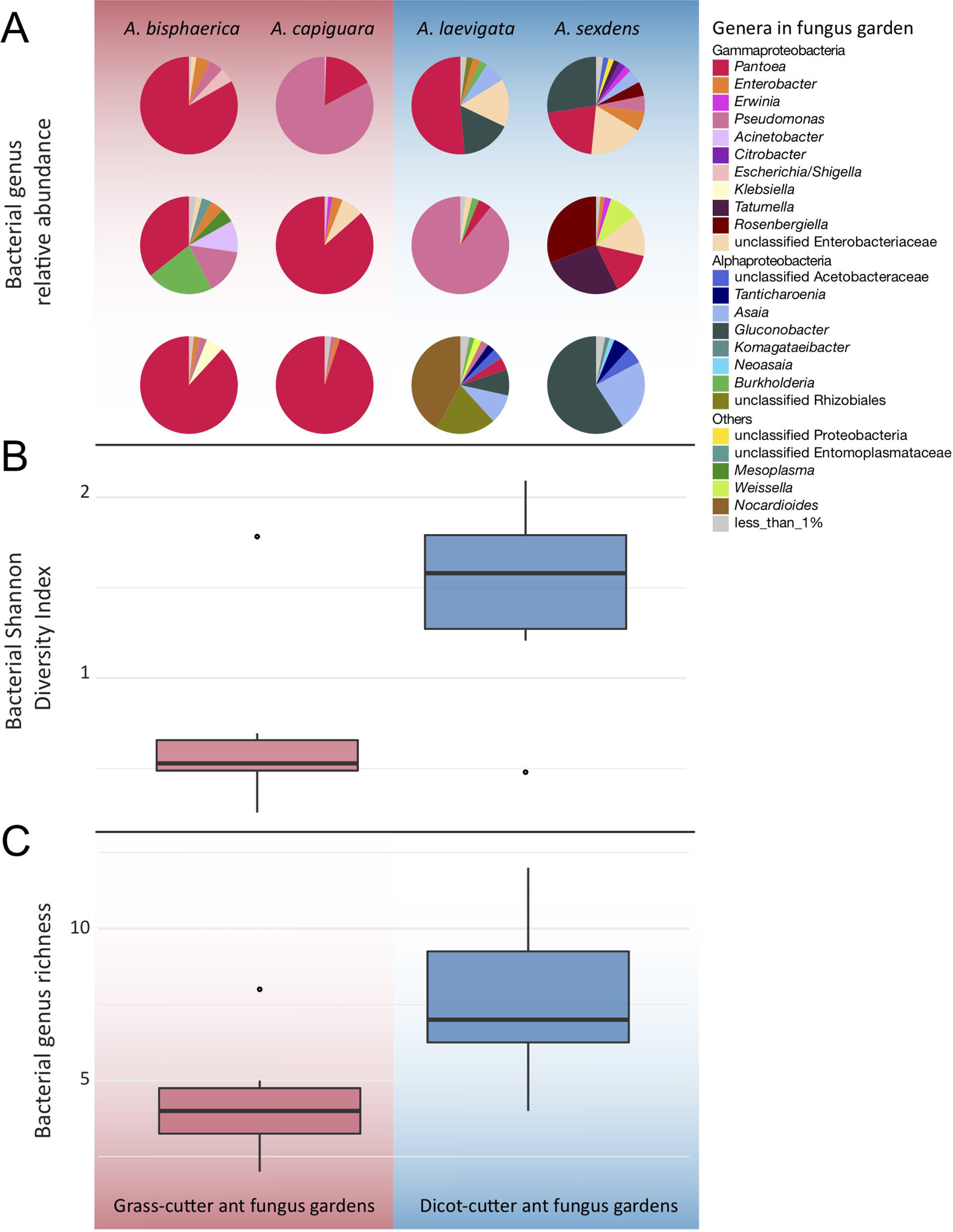
Genus-level bacterial community analysis of leaf-cutter ant fungus gardens from grass- and dicot-cutter ants, demonstrating that dicot-cutter ant fungus gardens have a higher diversity of bacteria. All data are based on 16S sequences extracted from the metagenomes using MATAM, and are relative abundances. A. Pie charts showing proportions of different bacterial genera in the fungus gardens B. Shannon diversity index of bacterial genera (for genera that consist of more than 1% of the total normalized gene count). C. Bacterial genus richness (for genera that consist of more than 1% of the total normalized gene count).

### Bacterial functional analysis

Overall, we found significant differences in the predicted bacterial community functional profiles between grass- and dicot-cutter ant fungus gardens (Figure 4). All individual bacterial genes that were significantly different between grass- and dicot-cutter ant fungus gardens are listed in Supplemental Table 1. In total, 514 predicted bacterial genes were significantly enriched, with 313 and 201 genes significantly enriched in grass- and dicot-cutter ant gardens, respectively (Supplemental Table 2, Supplemental Figures 4-6). Grass-cutter ant fungus gardens were enriched for amino acid biosynthesis genes for phenylalanine, tryptophan, tyrosine, histidine, arginine, lysine, cysteine, methionine, glycine, serine and threonine. They were also significantly enriched in terpenoid and siderophore biosynthesis genes (Figure 5) and had a significantly higher relative abundance of a gene in the nitrogen fixation pathway, nitrogenase molybdenum-iron protein beta chain (Supplemental Table 2). Dicot-cutter ant fungus gardens were particularly enriched in membrane transport genes (Figure 5).

**Figure 4.**
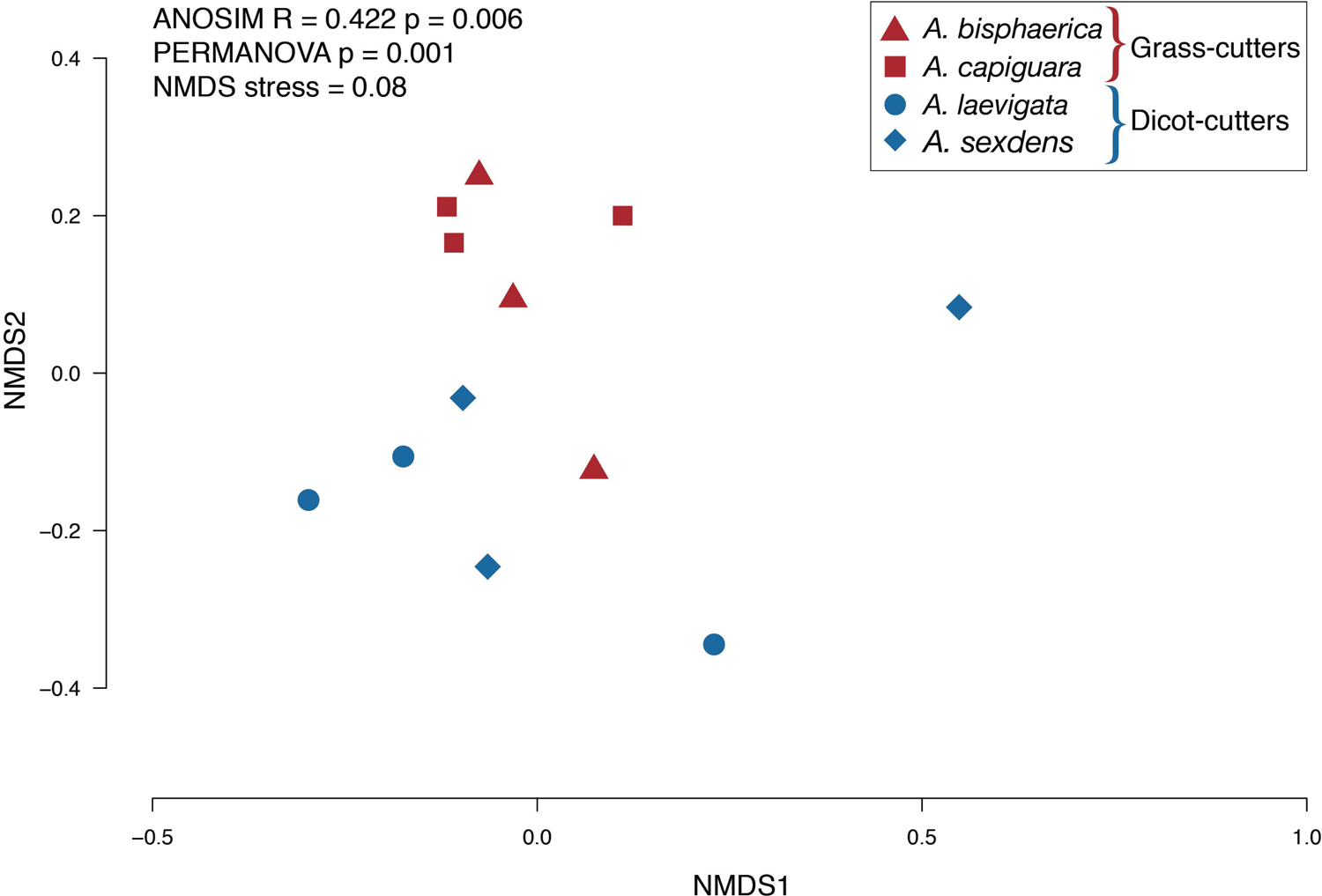
NMDS plot of KO functional genes from grass- and dicot-cutter ant fungus gardens. The KO profiles are significantly different between the fungus gardens of ants cutting the different substrates.

**Figure 5.**
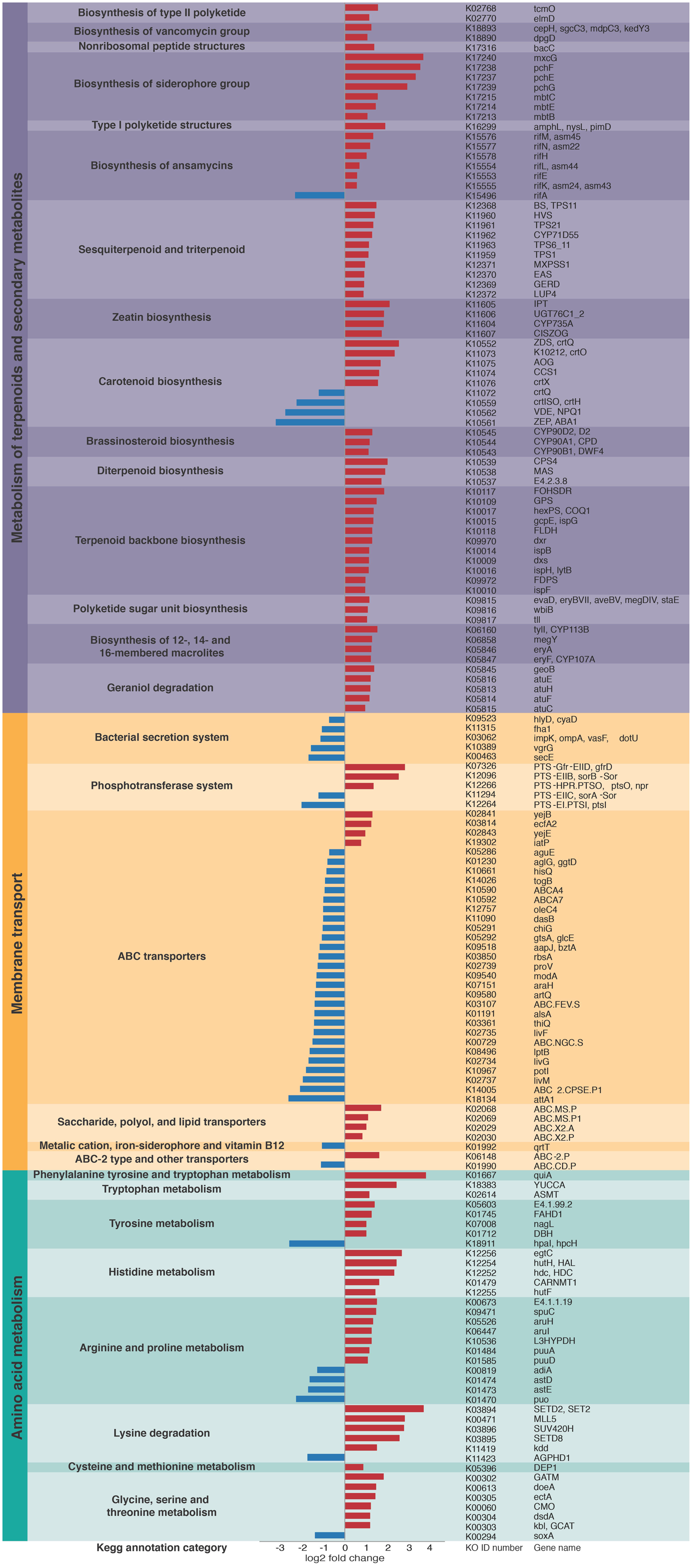
Particular groups of genes are enriched in either the grass- or dicot-cutter ant fungus gardens. Grass-cutter ant fungus gardens are enriched for genes involved in metabolism of terpenoids and other secondary metabolites, as well as genes involved in amino acid metabolism. In contrast, dicot-cutter ant fungus gardens are enriched for genes involved in membrane transport. Bars extending to the left (blue) represent genes that are significantly more abundant in dicot-cutter ant fungus gardens and bars extending to the right (red) represent genes that are significantly more abundant in grass-cutter ant fungus gardens.

### Plant taxonomy and consistency

The incorporated plant material was markedly different in consistency between the fungus gardens. *Atta bisphaerica* and *A. capiguara* gardens both contained material that was clearly grass, which was not triturated (Figure 1). In contrast, the leaf material in the fungus gardens of *A. laevigata* and *A. sexdens* was triturated to the point of being unrecognizable as plant material (Figure 1). We detected 68 plant species based on the *MatK* gene query in the metagenomes, from 40 genera and 15 families (Table 3). The fungus gardens of dicot-cutter ants had a significantly higher richness of plant genera than those of grass-cutter ants (ANOVA F=9.14, p=0.0128). As expected, the grass-cutter ant fungus gardens all contained grass (*Paspalum,* Poaceae). The dicot-cutter ant fungus gardens contained more genera and families of plants, which were mostly dicots, but three of these fungus gardens also contained some grass (Table 3). Although *A. laevigata* has been described in the literature as a grass and dicot-cutter, in this work we consider it a dicot-cutter for several reasons: the fungus garden consistency and amount of leaf trituration were like those of other dicot-cutter ants (Figure 1), the *MatK* data show that grass was not found in the fungus gardens of *A. laevigata* colony 2 (Table 3), and finally, the colonies used in the study were observed to be cutting mostly dicots (personal observation). As a result, we would expect that the bacterial community would be exposed to conditions most similar to other dicot-cutter ant fungus gardens.

**Table 3.**
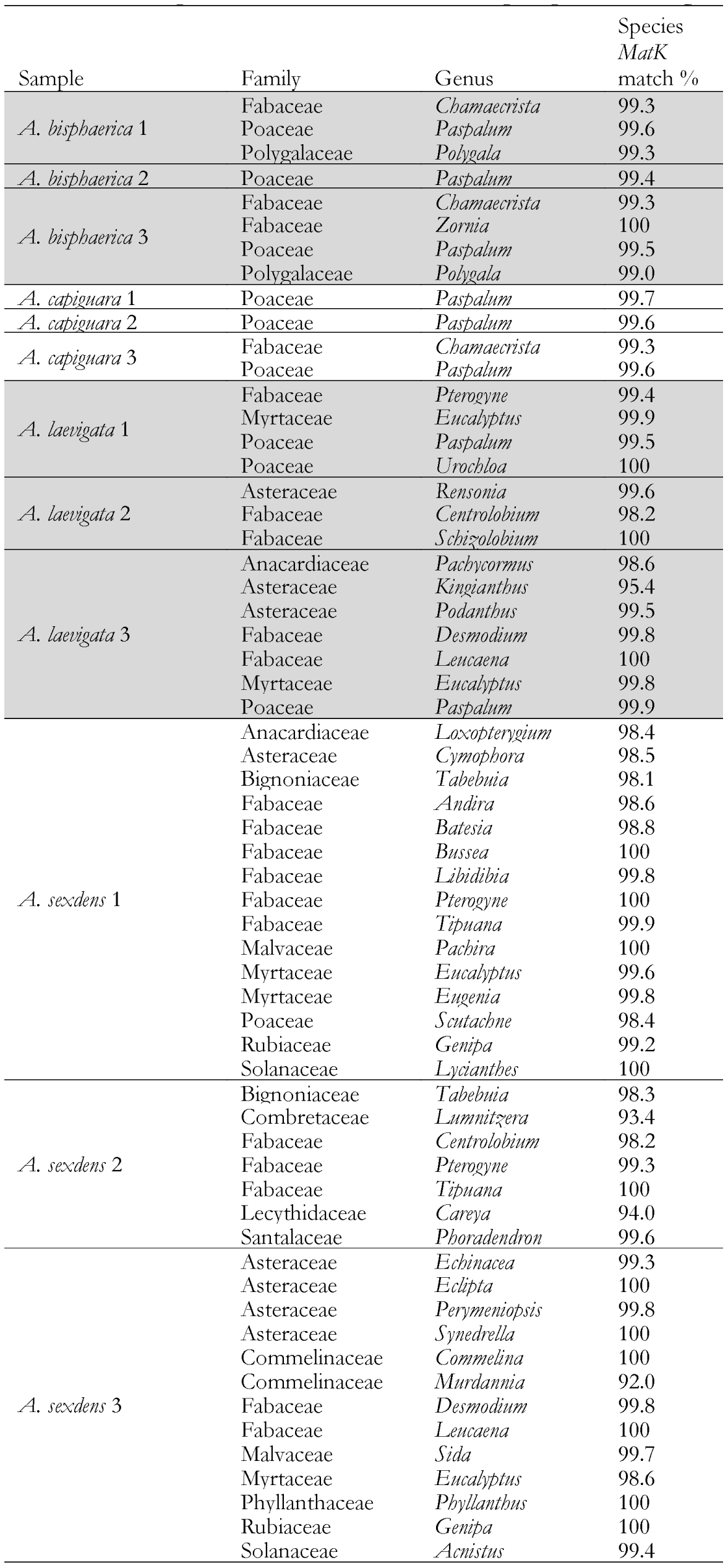
Plant genera detected in each fungus garden sample using *MatK* gene

### Iron content

The iron content of the fungus gardens is displayed in Figure 6, as mg of iron per kg of biomass. This represents the iron in the fungus garden, contained within both the plant and fungus material, but originating in the plants. The grass-cutter ant fungus gardens have lower amounts of iron than the dicot-cutter ant fungus, but this difference is not significant due to the high variability between *A. sexdens* gardens.

**Figure 6.**
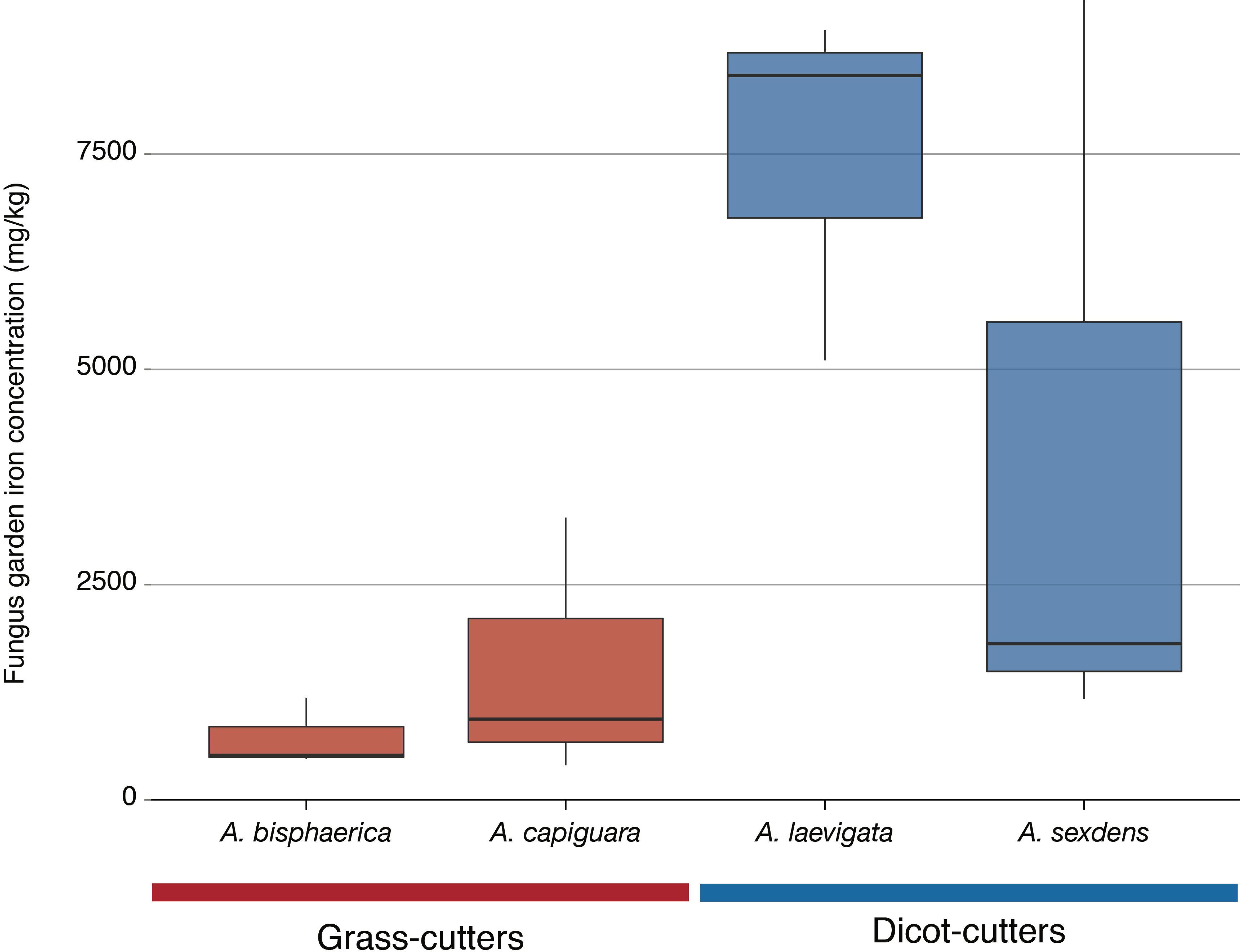
Iron content of fungus gardens from this study as measured by inductively coupled plasma optical emission spectroscopy. The iron content in the grass-cutter ant fungus gardens was lower than in the dicot-cutter ant fungus gardens. This difference is not statistically significant, however, since the *A. sexdens* fungus garden iron content is highly variable.

## Discussion

We can better understand animal diet specialization and transitions to novel substrates by understanding how microbial symbiont communities change in relation to transitions in host substrate use. *Atta* ants provide a relatively unique opportunity to examine a group of closely related herbivores that have transitioned from specialization on dicots to grasses. Dicots and grasses differ in terms of their cell wall composition, nutrient density and plant defense compounds. Here, using metagenomic sequencing, we examine this transition in the bacterial community in the fungus gardens of ants that are specialized on these different substrates. The results of this study demonstrate that the bacterial community differs in composition and functional capacity, depending on the type of substrate the ants incorporate into their gardens. These differences suggest that the bacteria might play a role in mediating the relationship between leaf-cutter ant colonies and the plants they consume.

If bacteria in fungus gardens are responsible for the breakdown of recalcitrant plant biomass, which is found in plant cell walls, we expect that the bacterial communities in the two ant groups examined here would be differentially enriched in the genes necessary for plant biomass breakdown. Grasses have a unique cell wall structure, containing (1→3),(1→4)-β-D-glucan chains and silica, neither of which are present in dicots (Popper and Tuohy, 2010). In other systems specialized on grass biomass breakdown, the microbes responsible for this produce specialized enzymes (King *et al.*, 2011) and have genomes that are adapted for this function (Wolfe *et al.*, 2012). In this system, between the bacterial communities in grass- and dicot-cutter ant fungus gardens, we do not identify a significant difference in the relative abundance of genes responsible for plant biomass degradation, namely glycoside hydrolases, carbohydrate esterases, carbohydrate binding molecules, polysaccharide lyases, and glycosyl transferases (Supplemental Table 1). Thus, we can conclude that garden bacteria do not respond to changes in cell wall structure between grasses and dicots. Instead, it is likely that the genome or gene expression in the fungus from these two systems would show differences, especially since the fungus is the primary degrader of plant biomass in leaf-cutter ant fungus gardens (Aylward *et al.*, 2013; Khadempour *et al.*, 2016; Nagamoto *et al.*, 2011; Grell *et al.*, 2013).

Leaf-cutter ants, in general, cut an exceptionally broad diversity of plants (Solomon, 2007; Mayhé-Nunes and Jaffe, 1998) and thus, have the potential to encounter a myriad of plant defense compounds that are toxic to themselves and their fungal cultivar. The ants are not enriched in gene families for plant defense compound detoxification (Rane *et al.*, 2016), so they must reduce the intake of these chemicals in other ways. Plant defense compound avoidance occurs in several steps. First, ants avoid cutting plants that contain plant defense compounds that are particularly toxic or abundant (Wirth *et al.*, 1997; Hubbell *et al.*, 1984; Howard, 1988). Second, many plant defense compounds that the ants encounter are volatile chemicals (Howard, 1988; Howard *et al.*, 1988), and in the time that the ants cut and carry the leaf material back to their colonies, some of the volatiles will have had time to dissipate. Finally, ants often leave leaf material in caches before they incorporate them into their fungus gardens (Hart and Ratnieks, 2000; Roschard and Roces, 2003), providing further opportunity for the defense compounds to evaporate. Nevertheless, some amount of volatiles can make their way into the gardens. In this study, using gas chromatography, we were able to detect eucalyptus-related compounds (eucalyptol, α-pinene, β□:pinene, *p*-cymene and *γ*-terpinene) in the fungus garden of one ant colony (*A. laevigata* 1) that was observed cutting considerable amounts of eucalyptus (Supplemental methods and Supplemental Figure 2).

In order to mitigate the deleterious effects of plant defense compounds, we expect the fungal cultivar *L. gongylophorus* would produce enzymes to degrade them. Indeed, work by De Fine Licht *et al.* (2013) implicates an important role for laccases from the fungal cultivar in detoxifying plant defense compounds. Nevertheless, bacteria in the garden may also play a role in mediating plant defense compounds, especially those that are toxic to the fungus, or that the fungus does not have the capacity to detoxify. Some evidence that bacteria found in dicot-cutter ant fungus gardens are better equipped to contend with toxic plant compounds is the higher abundance of membrane transport genes, especially ABC transporters in dicot-cutter ant fungus gardens (Figure 5), as these are known to be important in responding to toxins (Putman *et al.*, 2000; Glavinas *et al.*, 2004). The bacterial community contains the genes necessary for plant defense compound remediation, including many cytochrome P450s, gluthione S-transferases, and other genes involved in xenobiotic degradation (Supplemental Table 1), and aromatic compound degradation (Supplemental Figures 4 and 5), but they are not consistently enriched in the dicot-cutter ant fungus gardens. We expected that since dicot-cutter ants incorporate a higher diversity of plants into their gardens (Table 3), that the diversity of bacteria would also be higher in these gardens, and that the bacteria would have a higher capacity for the degradation of these defense compounds. While we did observe a greater diversity of bacteria in the dicot-cutter ant fungus gardens (Figure 3) we did not see a significant enrichment of plant defense compound degradation genes in these gardens (Figure 5, Supplemental Table 1). However, we still cannot exclude the possibility that bacteria are taking part in this process. Since each dicot-cutter ant colony cuts a unique set of plants (Table 3), they potentially encounter a unique set of plant defense compounds. If the bacterial community were to respond in a substrate-specific manner to different plant defense compounds, our analysis in this study would not reveal that. To elucidate the role of bacteria in plant defense compound remediation, closely controlled experiments with particular defense compounds of interest applied to bacterial cultures and to fungus gardens would be necessary.

Pinto-Tomas *et al.* (2009) established that *Pantoea* and *Klebsiella* bacteria in Central American leaf-cutter ant fungus gardens are supplementing the ant diet through nitrogen fixation. Plant material, in general, is low in nitrogen, and many herbivores supplement their diets through bacterial nitrogen fixation (Douglas, 2009; Hansen and Moran, 2013). Grasses are especially low in nitrogen (Mattson, 1980; Winkler and Herbst, 2004), so we predict that grass-cutter ant fungus gardens would be enriched in nitrogen-fixing bacteria with a corresponding enrichment of nitrogen-fixing genes. Here we show that a nitrogenase molybdenum-iron protein beta chain gene is significantly more abundant in grass-cutter ant fungus gardens (Supplemental Table 1). Other genes that are related to nutrient acquisition are also significantly more abundant in the grass-cutter ant fungus gardens (Figure 5), such as genes in amino acid metabolism pathways. While it has been shown that nitrogen fixed by bacteria is incorporated into the bodies of ants (Pinto-Tomás *et al.*, 2009), animals cannot simply absorb nitrogen as ammonium or nitrate, they require it to either be in the form of amino acids or other organic nitrogen-containing compounds (White, 1993). The enrichment of arginine biosynthesis genes is of particular interest since the genome of *Atta* is deficient in genes in this pathway (Suen *et al.*, 2011b). While a transcriptome study of *L. gongylophorus* demonstrated that the cultivar has the genes necessary for arginine biosynthesis (De Fine Licht *et al.*, 2014), the bacteria could supplement this process.

Other categories of genes enriched in the grass-cutter ant fungus garden bacteria are those involved in metabolism of terpenoids and other secondary metabolites, especially their biosynthesis. Grass-cutter ant fungus gardens are significantly enriched in 67 of these genes. This list includes seven siderophores, which are responsible for iron acquisition (Crosa, 1989; Winkelmann, 2002). Siderophores are costly to produce so the enrichment of these genes suggests that iron acquisition is important in this system. The grass-cutter ant fungus gardens examined in this study contained lower amounts of iron than the dicot-cutter ant fungus gardens (Figure 6). Since iron is an important cofactor in cells, and typically has low concentration in soils, it is a limiting resource for plants and the organisms that feed on them and in plant cells, it is found primarily in the cytoplasm (Briat *et al.*, 2006). We consider the lower iron content a proxy for plant nutritional density. In this system, bacteria that can produce more siderophores would be selected for over other bacteria that cannot produce as much, especially in the relatively low-iron grass gardens. The iron acquired by these bacteria could then potentially be used by the fungus or ants or they could remain in the bacteria. The movement of iron in the leaf-cutter ant fungus gardens provides an interesting and unexplored avenue for future studies.

Terpenoids are the most abundant secondary metabolites found in plants, and serve diverse roles (Langenheim, 1994; Gershenzon and Dudareva, 2007). The majority of research into the connection between plant terpenoids and animal-microbe symbioses are in regards to the detoxification of terpenes that would be deleterious to the animal host (Cheng *et al.*, 2013; Raffa, 2013; Adams *et al.*, 2013; Boone *et al.*, 2013; Wang *et al.*, 2012). However, not all terpenes are toxic to all organisms (Raffa, 2013), and in at least one instance they have been shown to supplement a herbivore’s diet after some modification by a gut bacterium (Berasategui *et al.*, 2017). Dicots contain higher quantities of terpenoids (Wetterer, 1994; Mariaca *et al.*, 1997). One possibility is that the bacteria in these fungus gardens are producing terpenes as a nutritional additive, especially in the grass-cutter ant fungus gardens where there are lower terpene inputs and these genes are enriched (Figure 5, Supplemental Figure 2).

Overall, this study demonstrates that the bacterial community differs in both community composition and functional capacity between grass- and dicot-cutter ant fungus gardens. We argue that the difference in functional capacity of bacteria in the different gardens can be used by the fungus and, downstream of that, the ants in this system. As an extension of this, the bacteria may play a role in allowing grass-cutter ants to utilize a lower quality substrate than their dicot-cutter counterparts. Of course, this is all based on metagenomic work, and has its limitations. First, the steps of transcription and translation lie between the number of gene copies and ultimate function. However, previous efforts to examine the bacterial community *in situ*, both through metaproteomics and metatranscriptomics, have not been fruitful, since the fungal biomass, transcripts, and proteins swamp out the bacterial portion (Khadempour *et al.*, 2016; Moreira-Soto, 2016). Therefore, to date, this work provides the most comprehensive picture of the bacterial community in leaf-cutter ant fungus gardens and the potential roles they may play in the system.

A second limitation is that we only present relative abundances, rather than absolute, and therefore assume that the overall proportion of bacteria to other biomass in the fungus gardens is consistent, which is especially important since bacteria represent such a small proportion of the overall biomass in the gardens. It might be argued that differences in statistical abundance will not be reflected in biological differences, or that the abundance of the bacteria is too low overall to have any effect on the system as a whole. However, previous work demonstrated that even with their low absolute abundance in leaf-cutter ant fungus gardens, the nitrogen that is fixed by bacteria is taken up by the ants (Pinto-Tomás *et al.*, 2009). The impact of a low-abundance community member has also been shown in another microbiome (Romano *et al.*, 2015). So we believe that the differences we observe can have impacts on the system.

With the limitations described above, we cannot definitively conclude that fungus garden bacteria expand the diversity of plants that ants can incorporate into their gardens, but we have opened up several new avenues of research. Follow-up studies will focus on bacteria both in culture and in fungus gardens using more targeted approaches to either confirm or contradict the results of this study, which suggest that the bacteria in leaf-cutter ant fungus gardens may play a role in mediating the relationship between ants and the types of plants that they incorporate into their gardens.

## Acknowledgements

The authors would like to thank Andre Rodrigues of UNESP for his help with collections and permits. This work was funded in part by the U.S. Department of Energy Great Lakes Bioenergy Research Center (DOE Office of Science BER DE-FC02-07ER64494), National Institute of Food and Agriculture, United States Department of Agriculture, under ID number 1003779, National Institutes of Health (NIH grant U19TW009872) and São Paulo Research Foundation (FAPESP grant 2013/50954-0). Collection at the UNESP campus in Botucatu was completed under the ICMBio permit 31534-1. Collection at the USP campus in Riberão Preto was completed under the SISBIO permit 46555-5 and CNPq permit 010936/2014-9. The work conducted by the DOE Joint Genome Institute, a DOE Office of Science User Facility, is supported by the Office of Science of the DOE under Contract No. DE-AC02-05CH11231.

