## Supplementary figures and images for "Metagenomics reveals diet-specific specialization of bacterial communities in fungus gardens of grass- and dicot-cutter ants"

### Supplemental Figure 1

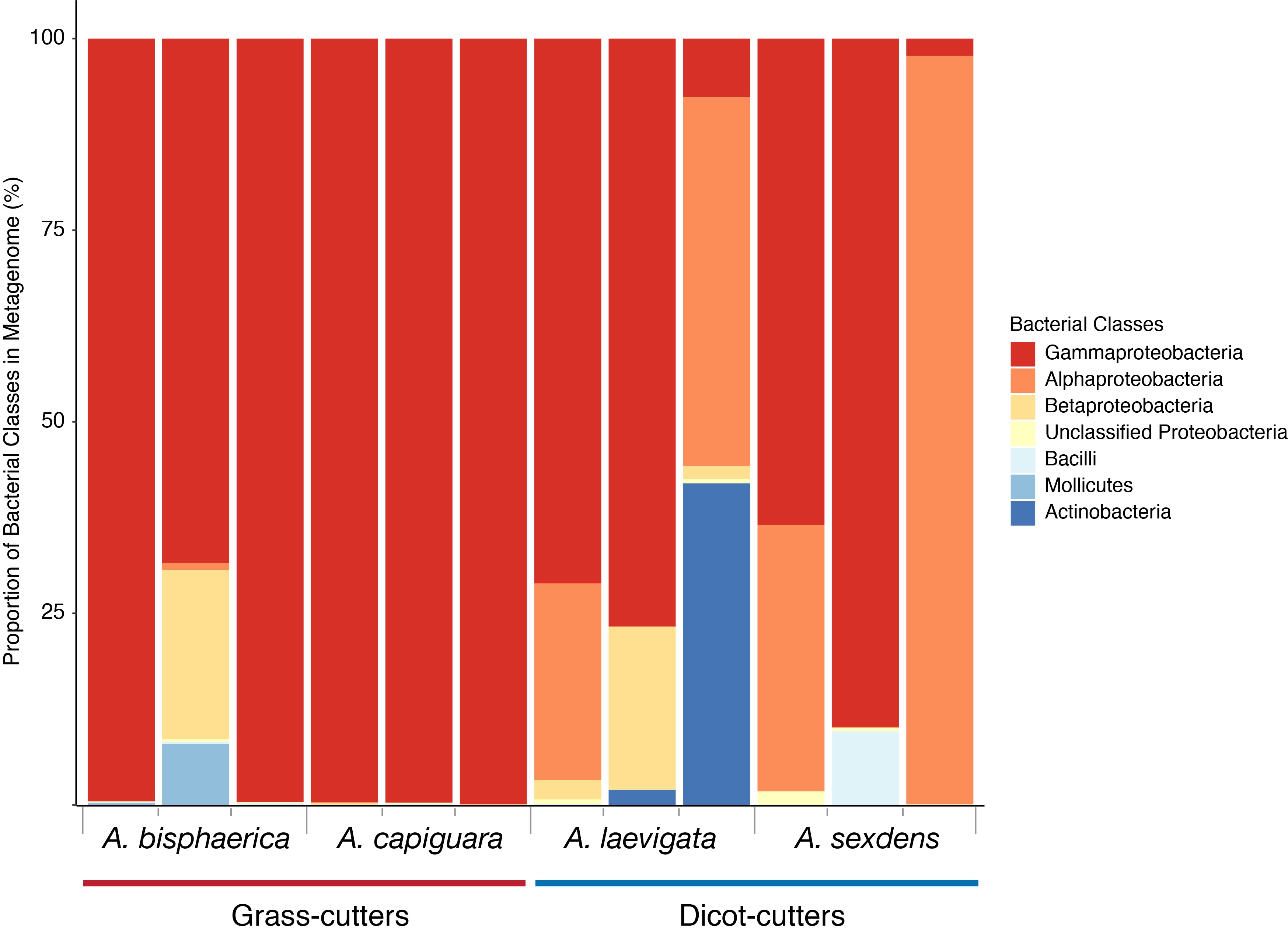

### Supplemental Figure 2

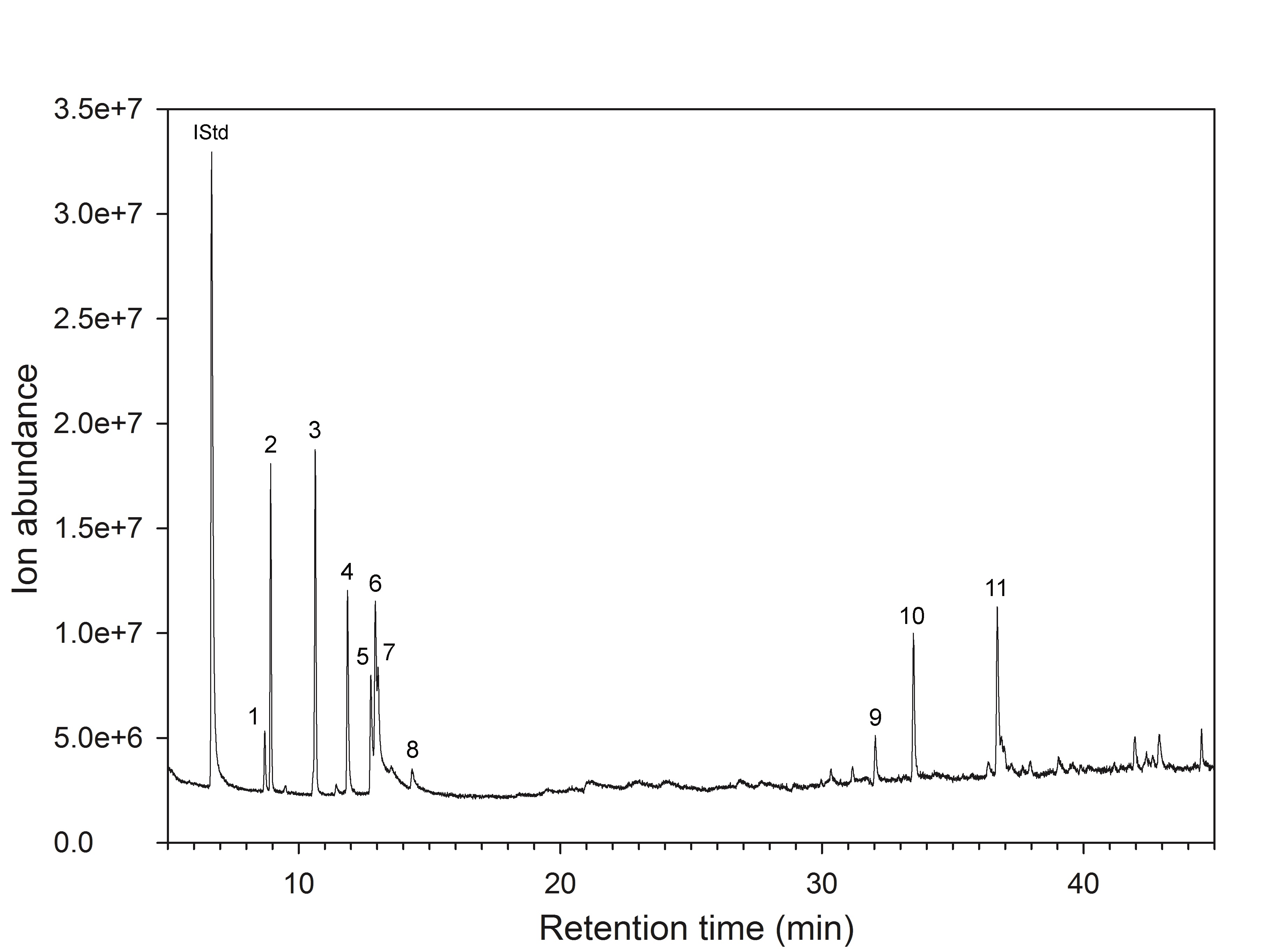

### Supplemental Figure 3

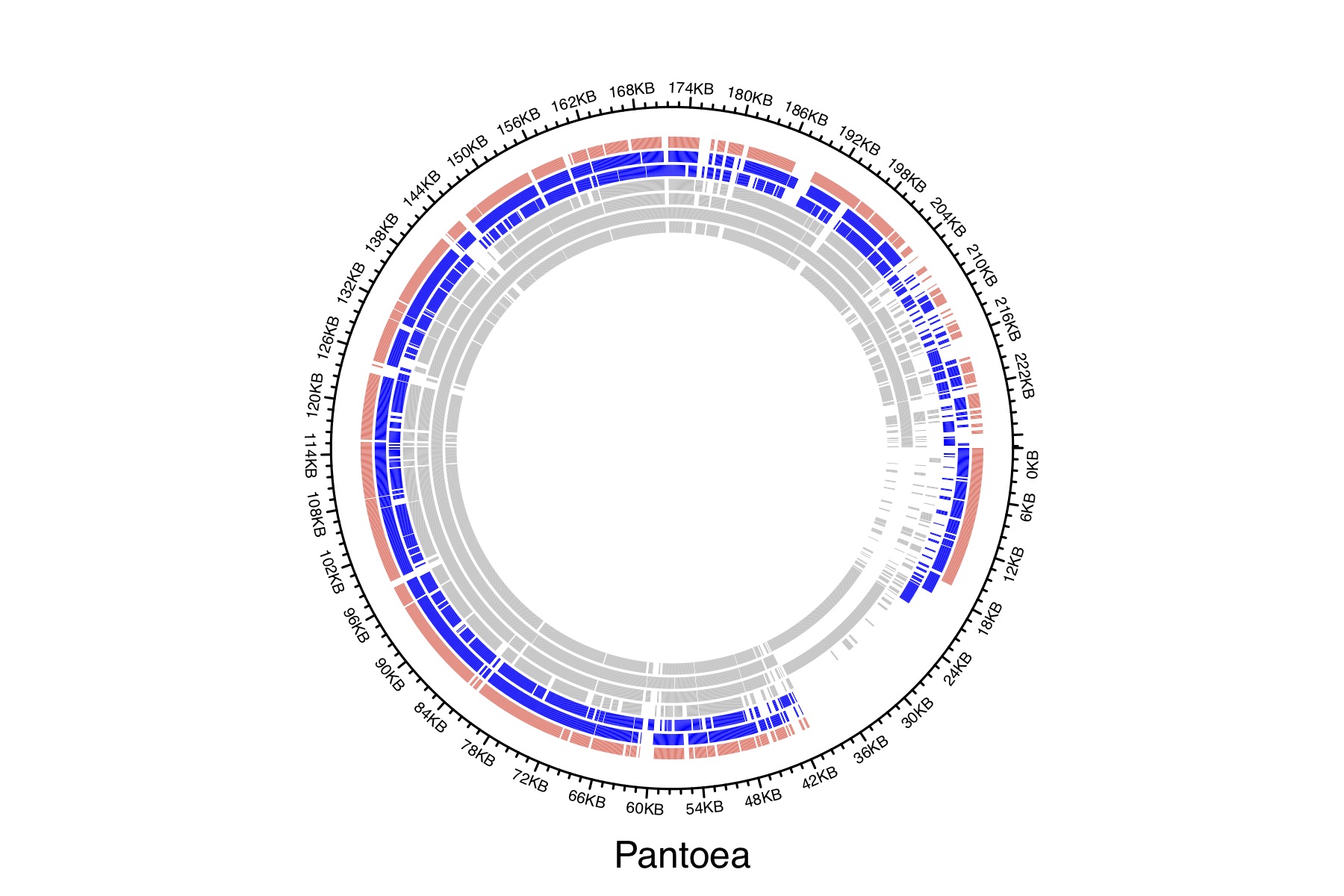

### Supplemental Figure 4

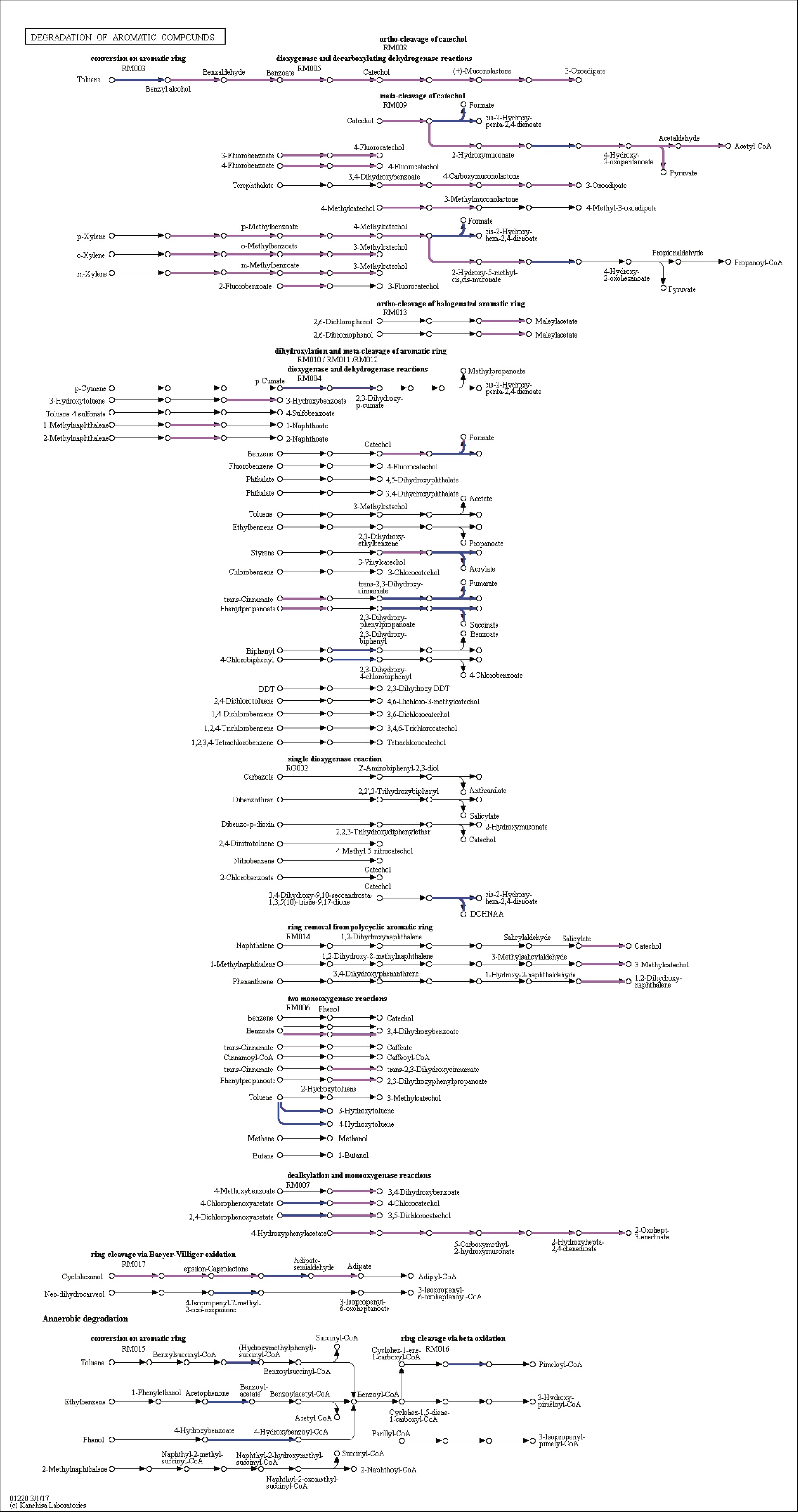

### Supplemental Figure 5

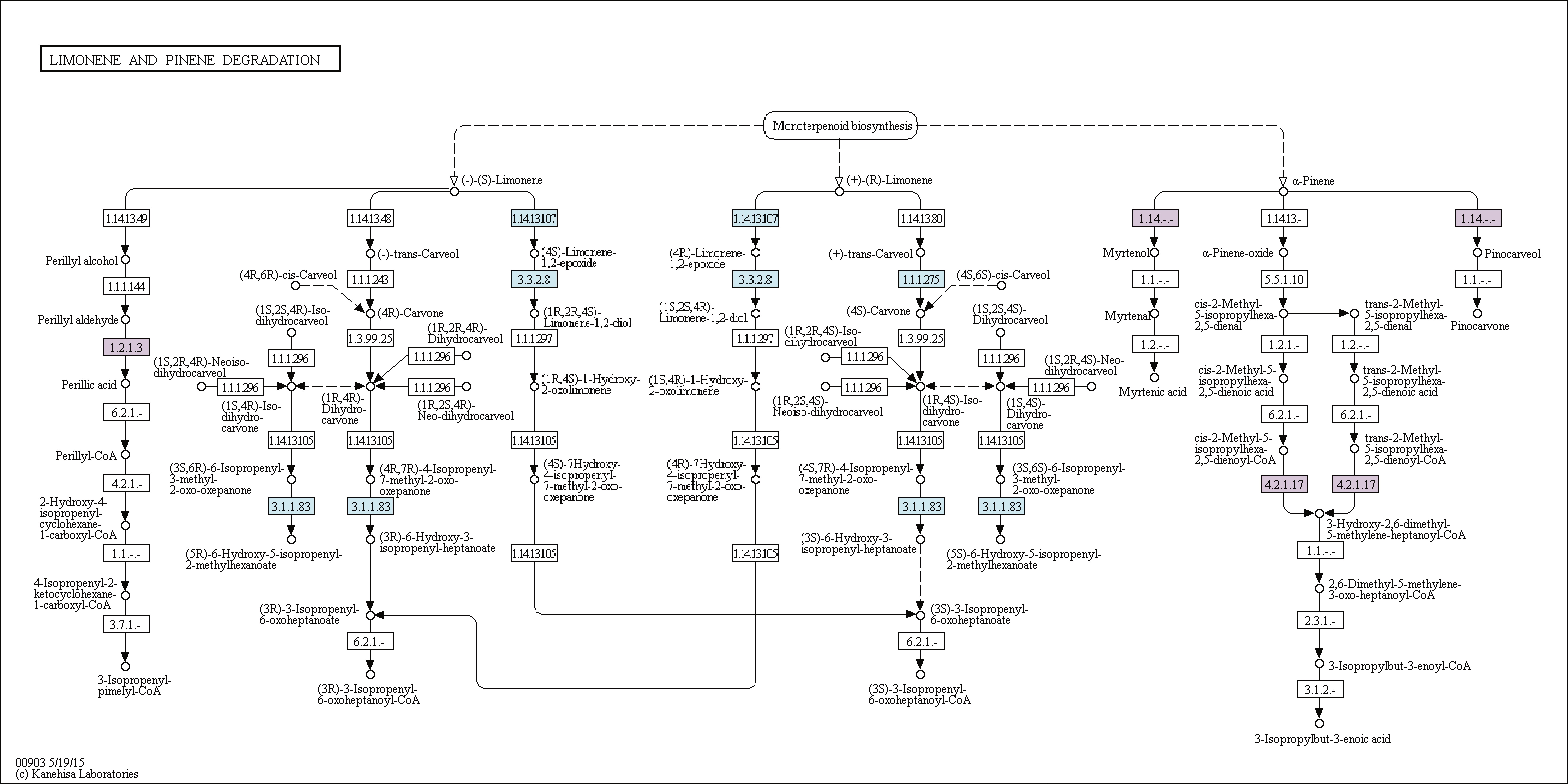

### Supplemental Figure 6

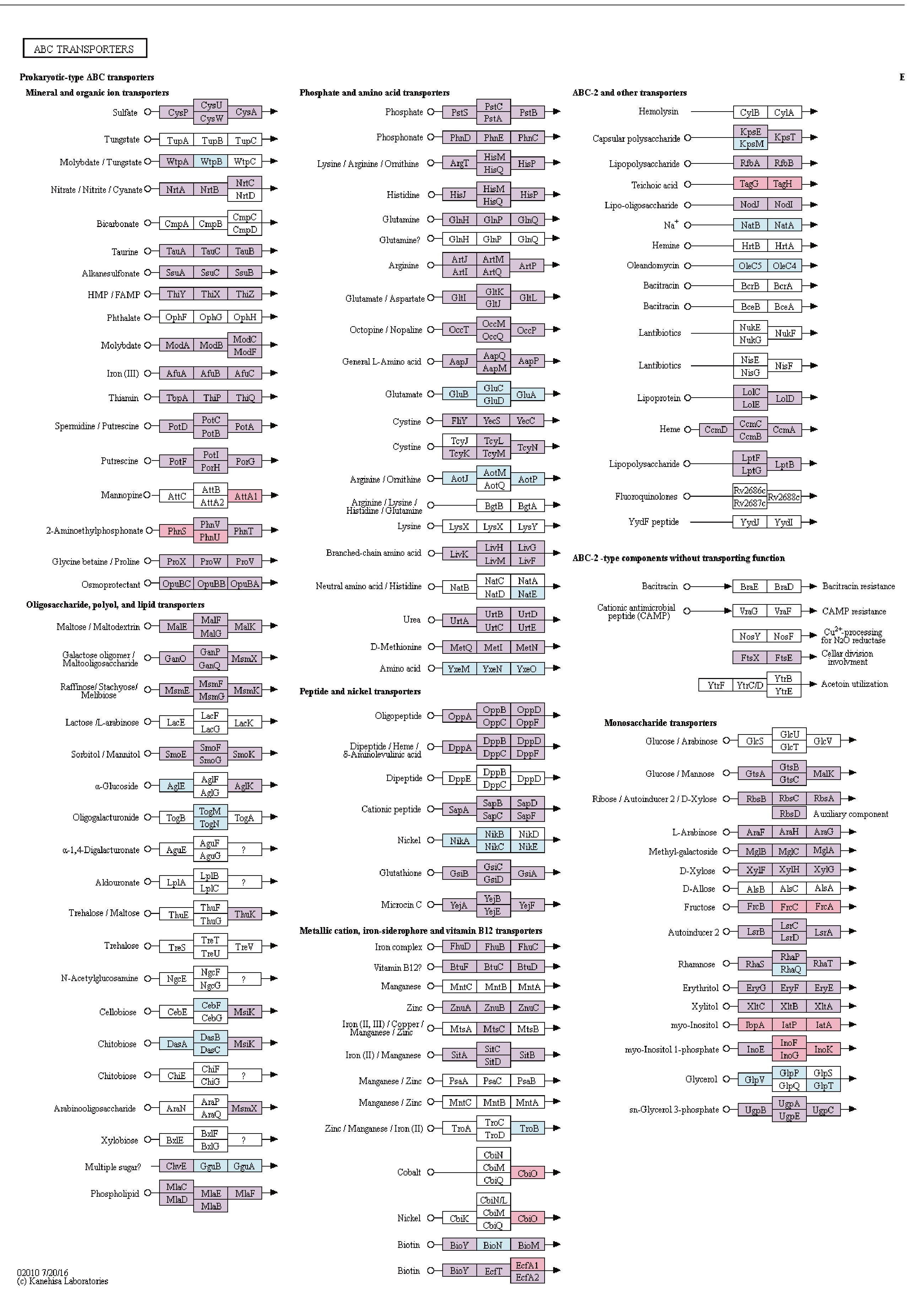
