## Supplemental Text for "Metagenomics reveals diet-specific specialization of bacterial communities in fungus gardens of grass- and dicot-cutter ants"

### Supplemental Materials

#### Supplemental Figure captions

Supplemental Figure 1 Bar plots showing the relative abundance of different bacterial classes, dominated by Gammaproteobacteria.

Supplemental Figure 2 GC-MS total ion chromatogram of terpenes extracted from *Atta laevigata* (1) fungus garden material. IStd = internal standard (*m*-xylene), 1 = α-thujene, 2 = α-pinene, 3 = β-pinene, 4 = α-phellandrene, 5 = *p*-cymeme, 6 = limonene, 7 = eucalyptol (1,8-cineole), 8 = γ-terpinene, 9 = alloaromadendrene, 10 = unknown sesquiterpene, 11 = caryophyllene oxide. Terpenes were identified by both their mass spectra and retention indices calculated with an *n*-alkane series. No other samples presented identifiable peaks.

Supplemental Figure 3 Pangenome of *Pantoea* composite genomes. The outermost band is a count of all kilobases represented. The next seven rings represent *Pantoea* genomes. The red circle is from a grass-cutter ant fungus garden (ACBM2) the two blue genomes are from dicot-cutter ant fungus gardens (ALBM2 and ASBM1) and the next four grey genomes are from isolates of *Pantoea* (Accession numbers: SAMN08357569, SAMN08357571, SAMN08357570, and SAMN00002769). From the limited composite genomes we compiled, we did not identify patterns in gene content dependent on ant substrate specialization.

Supplemental Figure 4 KEGG pathways involved in the degradation of aromatic compounds. Blue arrows indicate genes that are present in dicot-cutter ant fungus gardens, red arrows indicate genes that are present in grass-cutter ant fungus gardens and purple arrows indicate genes that are present in both.

Supplemental Figure 5 KEGG pathways involved limonene and pinene degradation. Blue boxes indicate genes that are present in dicot-cutter ant fungus gardens, red boxes indicate genes that are present in grass-cutter ant fungus gardens and purple boxes indicate genes that are present in both.

Supplemental Figure 6 KEGG pathways involved in ABC transporters. Blue boxes indicate genes that are present in dicot-cutter ant fungus gardens, red boxes indicate genes that are present in grass-cutter ant fungus gardens and purple boxes indicate genes that are present in both.

#### Supplemental methods

##### Composite genomes

Composite genomes were created using the MaxBin program with default settings (Wu *et al.*, 2014). Their quality, contamination and completeness were assessed using CheckM (Parks *et al.*, 2015). Only composite genomes with less than 10% contamination and more than 90% completeness were included in further analysis. CheckM reports the taxonomy level of each composite genome based on the most recent common ancestor of the annotation of the core genes identified, thus not all of them were identified to an informative level. To further elucidate the taxonomy of each composite genome, they were annotated separately using Prokka (Seemann, 2014). If a 16S sequence was recovered from a composite genome, then that sequence was blasted in NCBI. The only group that we had clear taxonomic identification, and which was present in both grass- and dicot-cutter ant fungus gardens, was *Pantoea*. Pangenomic analysis was carried out including three composite *Pantoea* genomes and four published *Pantoea* genomes using GET_HOMOLOGUES (Contreras-Moreira and Vinuesa, 2013). 2450 genes were shared at least by two out of those seven genomes. Circle plots showing the presence and absence of those 2450 genes in each genome were created using the Circlize package (Gu *et al.*, 2014) in R (R Core Team, 2013).

##### KEGG orthology pathways

To map out the completeness of various KEGG pathways, first presence/absence was determined for each KEGG gene. A gene was considered present in either grass or dicot-cutter ant fungus gardens if it was present in at least 4 of the 6 metagenomes in that category. Presence and absence in grass-cutter ant fungus gardens, dicot-cutter ant fungus gardens or both were plotted on KO pathways through the Kegg Orthology Database website (http://www.genome.jp/kegg/ko.html). The pathways are represented in Supplemental Figures 4-6.

##### KEGG orthology annotations ascribed to genera

To determine which genera of bacteria were contributing to the abundance of particular functional genes, all genes matching to the most abundant genera (*Pantoea, Pseudomonas, Burkholderia, Enterobacter, Klebsiella,* and *Serratia*) were binned by genus. Next, the KEGG annotation was found for each gene and the numbers of genes matching to that annotation were added for each genus. Non-normalized and normalized gene counts for each genus are presented in Supplemental Table 2 in two separate tabs.

##### Gas chromatography

Fungus samples were collected from each of the twelve leaf-cutter ant colonies, then dried for 24 hours at 105°C. Samples were then crushed by mortar and pestle with liquid nitrogen. 100 mg of crushed material from each sample was weighed out and placed into 2.0 mL self-standing graduated microcentrifuge tubes.

To extract the compounds from the samples, 750 µl of methanol was added to each of the 100 mg of frozen powdered material. Each sample was briefly vortex mixed, sonicated in a chilled water bath for ten minutes, vortex mixed again, then centrifuged at 20,000 rpm for ten minutes. For the analysis of the volatile compounds, 1 μL of methanol extract from each the samples was placed individually into a coupled Trace 1310 gas chromatograph and Thermo ISQ mass spectrometer with electron ionization at 70 eV (GC-MS) machine with the injector temperature set to 260°C. Compounds were identified using both of their mass spectra compared to the NIST 2014 MS library, and using their linear retention indices calculated from separate injections of a hydrocarbon series (C_8_-C_20_).
